# Astrocytic Ca^2+^ signals partake in inhibitory neurovascular coupling in a brain state-dependent manner

**DOI:** 10.1101/2022.02.23.478942

**Authors:** Aske Krogsgaard, Leonora Sperling, Matilda Dahlqvist, Kirsten Thomsen, Gabriele Vydmantaite, Martin Lauritzen, Barbara Lykke Lind

## Abstract

Neurovascular coupling (NVC) modulates cerebral blood flow to match increased metabolic demand during neuronal excitation. Activation of inhibitory interneurons also increase blood flow, but the basis for this inhibitory NVC is unclear. We performed two-photon microscopy in awake mice to examine the correlation between astrocytic Ca^2+^ and NVC, evoked by activity in either all (VGAT_IN_) or parvalbumin-positive GABAergic interneurons (PV_IN_). Optogenetic stimulation of VGAT_IN_ and PV_IN_ in the somatosensory cortex triggered astrocytic Ca^2+^ increases that were abolished by anaesthesia. PV_IN_ evoked astrocytic Ca^2+^ responses with a short latency that preceded NVC, whereas VGAT_IN_ evoked Ca^2+^ increases that were delayed relative to the NVC response. The early onset in PV_IN_ evoked Ca^2+^ increases dependent on noradrenaline release from locus coeruleus, which also affected inhibitory NVC. Therefore, NVC mechanisms should be studied in awake mice and, though the relationship between interneuron activity and astrocytic Ca^2+^ is complex, we found a correlation between astrocyte activity and NVC in PV_IN_.

## Introduction

Astrocyte processes enwrap brain synapses as part of the tripartite synapse^1,2^. As such, they modulate synaptic strength, which has been studied extensively for excitatory glutamatergic signal transmission^3,4^. However, far less is known about how astrocytes respond to GABAergic inhibitory interneurons^5-8^.

Interneurons constitute a small fraction of cortical neurons but are critical in shaping network activity and signal integration^9^. Though inhibitory, interneuron activation can trigger neurovascular coupling (NVC)^10-12^, but the basis for this is unclear. Several different types of interneurons secrete vasoactive compounds^11-18^. However, some subtypes, such as the fast-spiking parvalbumin-positive GABAergic interneurons (PV_IN_), barely express nNOS^10^ or other vasodilators. Nevertheless, optogenetic studies in vivo indicate a PV_IN_ - induced increase in cerebral blood flow (CBF), which must then be mediated by a GABAergic effect^15,19^. Here, we hypothesize that GABA interacts with astrocytes to mediate PV_IN_-evoked NVC via release of astrocytic vasodilators^20-22^.

Astrocytes have been found to modulate NVC^23-26^, though the exact mode of action is still debated^27,28^. A proposed primary target for the regulation of NVC by astrocytes is the capillaries branching from penetrating arterioles^29,30^. Whether astrocytes contribute to the NVC evoked by interneurons is unknown. An effect of GABA on astrocytes is expected because they express both GABA_A_ and GABA_B_ receptors, as well as GABA transporters. Disconcertingly, studies in brain slices and anaesthetized mice have shown that astrocytes respond to GABA release through a delayed or fluctuating increase in astrocytic Ca^2+5-8^. This may due to the anaesthesia-induced depression of spontaneous Ca^2+^ transients in cortical astrocytes^31^. Similarly, the release of neuromodulators is impaired in slices, which can be critical since astrocyte responses reflect noradrenaline (NA)^32,33^ and brain state levels^34,35^. Therefore, to fully elucidate the astrocytic response, it must be assessed under experimental conditions free from these constraints.

The objective of the present study was to examine the impact of the awake state on GABA-elicited astrocytic Ca^2+^ responses and whether astrocytes mediate inhibitory NVC in the cortex of awake mice. In anaesthetized mice we found no immediate astrocytic Ca^2+^ responses to a local puff of GABA. Instead, it triggered a slow and fluctuating activity. In contrast, optogenetic stimulation of PV_IN_ in awake mice evoked astrocytic Ca^2+^ responses with short latency dependent on preserved NA signalling rather than GABA_A_ or GABA_B_ receptor activity. Stimulation of PV_IN_ also triggered vasodilation of both penetrating PAs and capillaries. This effect was impaired by anaesthesia and reduced by zolpidem, an allosteric modulator of GABA_A_ receptors. Capillary dilation was also reduced by baclofen, an activator of the GABA_B_ receptor. Simultaneous optogenetic stimulation of all interneuron subtypes (VGAT_IN_) influenced both astrocytes and NVC responses differently than stimulation of PV_IN_ alone. This shows that, under some conditions, interneuronal release of vasoregulators may increase local blood flow independent of astrocytes, and that the involvement of astrocytes in GABA-dependent NVC may be specific to PV_IN_ activity. Our findings underline that astrocyte and interneuron interactions should be studied in awake conditions. Doing so, we found that astrocytes may partake in GABAergic processes on a timescale permissive for the regulation of inhibitory synapses as well as NVC.

## Results

### In anesthetized animals, astrocytes respond to GABA with delayed and fluctuating Ca^2+^ activity

First, we examined how astrocytes respond to local exposure to GABA via a small-volume puff injection^36^. Two-photon imaging was performed in anaesthetized wild-type mice in layer II/III of the somatosensory cortex (Figure 1A). To monitor Ca^2+^ transients, the astrocytes were transduced to express the Ca^2+^ indicator GCaMP6f under the GFAP promotor tethered to the cell membrane, which allowed the activity in small perisynaptic processes to be monitored^37^.

**Figure 1:**
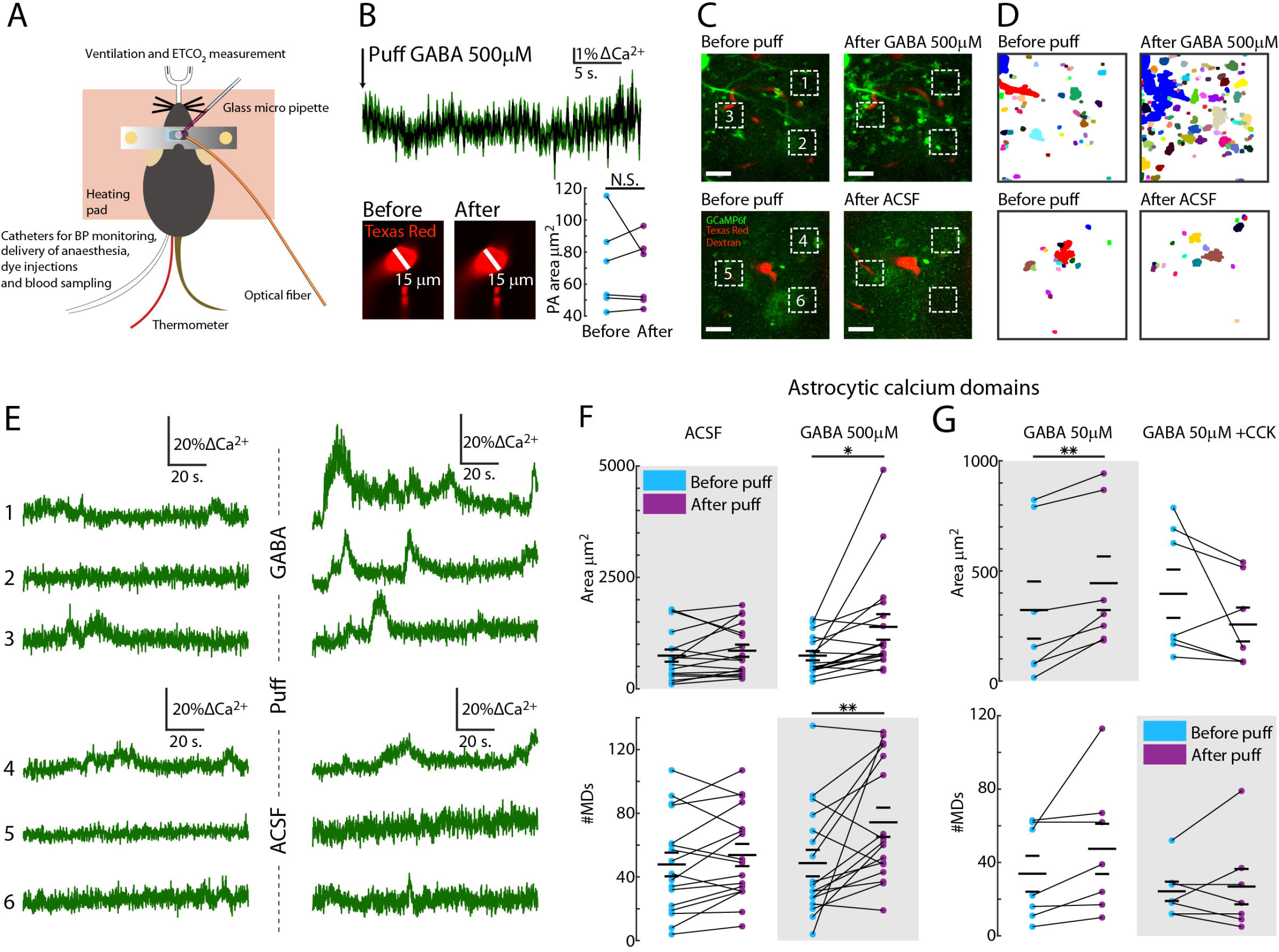
In anaesthetized animals, astrocytes show a delayed response to local GABA exposure, which is eliminated by the neuropeptide CCK. **A)** Schematic of the experimental setup in anaesthetized mice. GABA was introduced by puffing via a glass micropipette to study local effects. **B)** Upper panel: The data shows the average Ca^2+^ trace (± SEM in green) after the puff. Bottom panel: Corresponding example images of the penetrating arteriole (PA) before and after puffing. The Texas Red-filled micropipette used for puffing is visible at the bottom of the images. Right: The PA diameter and astrocytic Ca^2+^ levels were unaffected by a puff of 500 µM GABA (paired t-test, p=0.4165, n=6 mice). **C)** Maximum intensity projections of a field of view (FOV) before and after puffing 500 µM GABA or aCSF. Green is the astrocytic GCaMP6f signal, and red is the fluorescence of blood-circulating Texas Red dextran used to delineate the vasculature. Dashed boxes show the example areas with astrocytic Ca^2+^ analysed in (E). **D)** Map of active microdomains of astrocytic Ca^2+^ activity recognized by the automatic event detection algorithm. The output map shows the active areas before and after puffing GABA and aCSF. **E)** Astrocytic Ca^2+^ traces from the three ROIs indicated in (C) before and after puffing GABA (top three rows) and aCSF (bottom three rows). **F)** Area of active astrocytic Ca^2+^ microdomains (MDs) and the number of active astrocytic Ca^2+^ MDs were unchanged by puffing aCSF but increased after puffing 500 µM GABA. *p<0.05, **p<0.01, Student’s paired t-test (n=17 FOV, 12 mice). **G)** Top: Total area of active astrocytic Ca^2+^ MDs before and after puffing GABA on its own and GABA (50 µM) + CCK (25 nM). Bottom: Number of active astrocytic Ca^2+^ MDs detected before and after puffing GABA on its own (left) and GABA + CCK (right). **p<0.01, Student’s paired t-test (n=7 mice).

Local puff application of GABA did not have an immediate effect on astrocytic Ca^2+^ increase or change diameter of the penetrating arterioles (PA) (Figure 1B) but induced a delayed and fluctuating increase in Ca^2+^ (Figure 1C-1E). These astrocytic Ca^2+^ changes were quantified using an automatic event detection algorithm for unbiased identification of astrocytic Ca^2+^ increases^35,38^ (Figure 1D and Sup. Figure S1). To rule out the potential effects of mechanical stimulation by the pressure of the injectant, we compared the effect of GABA puffs to puffs with artificial cerebrospinal fluid (aCSF). Puffing 50 µM and 500 μM GABA induced prolonged increases in astrocytic Ca^2+^ (Figure 1F), measured as both the size of the active area (Student’s paired t-test, n=17 fields of view [FOVs] across 12 mice, p=0.016) and the number of active domains (n=17 FOVs, 12 mice, p=0.0091). In contrast, no increases were observed when puffing aCSF (n=17 FOVs, 12 mice, area p=0.2658 and number p=0.1238; Figure 1F).

As several neuropeptides are commonly co-released with GABA, we introduced cholecystokinin (CCK) into the puffing pipette with GABA to mimic stimulation of astrocytes by two different interneuron subtypes. This combination (25 nM CCK and 50 µM GABA) removed the prolonged fluctuations in astrocytic Ca^2+^ evoked by GABA alone (Figure 1G; all Student’s paired t-test; GABA only: area, n=7 mice, p=0.0024; number of MDs, n=7, p=0.1990; GABA + CCK: area, n=7, p=0.3582; number of MDs, n=7, p=0.4606). This reveals a complex astrocytic dependence on interneuron activity beyond the impact of GABA.

### Astrocytic Ca^2+^ responses to interneuron optogenetic activation are immediate, subtype-dependent, and diminished by anaesthesia

As the GABA effect on astrocytes seemed to be modified by co-released neuropeptides, we speculated that responses may depend on which interneuron subtype releases the GABA. Therefore, we used optogenetic stimulation (wavelength 473 nm, frequency 100 Hz, pulse duration 7.5 ms, pulse train duration 2 or 15 seconds) in two genetically modified mouse strains to further explore the influence of interneurons on astrocytic Ca^2+^. The first strain expressed channel rhodopsin (ChR2) in the PV_IN_, and in the second strain the VGAT promotor was used to express ChR2 in all GABAergic interneuron subtypes (VGAT_IN_). PV_IN_ are known to secrete only GABA^9^, whereas VGAT_IN_ stimulation is expected to release GABA along with several neuropeptides^10-12^. Using this paradigm, we could compare the effect of GABA alone to the combined influence of GABA and neuropeptides co-released by all interneuron subtypes. Optogenetic stimulation of both PV_IN_ and VGAT_IN_ inhibited the ongoing electrical activity (Figure 2B), indicating effective GABA release.

**Figure 2:**
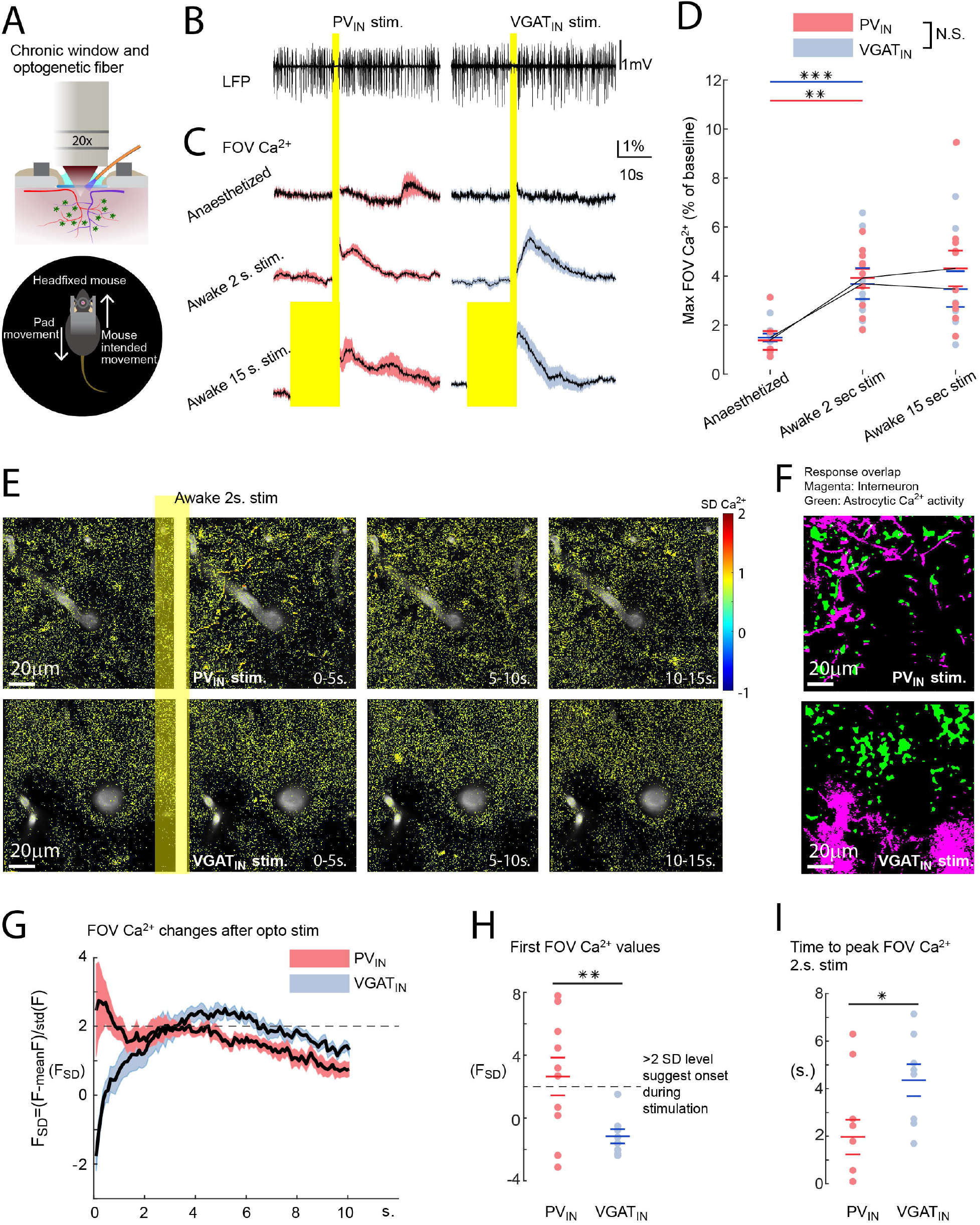
Optogenetic stimulation of interneurons triggers astrocytic Ca^2+^ increases in awake mice. **A)** Awake mice are free to move in the air-cushioned Mobile Neurotar HomeCage®, which combines stable head-fixation with a two-dimensional treadmill. **B)** Local field potential (LFP) responses clearly show the synaptic inhibition as dropouts of the potential during optogenetic stimulation of ChR2 in PV_IN_ and VGAT_IN_. The trace is an overlay from three repeated stimulations in one mouse. **C)** Average astrocytic field of view (FOV) Ca^2+^ responses to optogenetic ChR2 stimulation of PV_IN_ (red, n=10 FOV, 6 mice) and VGAT_IN_ (blue, n=8 FOV, 7 mice). From top: α-chloralose anaesthesia, awake mice 2-second, awake mice 15-second stimulation. Black lines indicate averages and shaded areas ± SEM. **D)** Maximal FOV Ca^2+^ response for the conditions in (C) show the effect of anaesthesia. The differences between the anaesthetized state and awake state for both strains were significant. **p<0.01, ***p<0.001. **E)** Ca^2+^ responses to optogenetic stimulation of ChR2 in PV_IN_ (Top) or VGAT_IN_ (Bottom) interneurons. Pseudocolours indicate normalized Ca^2+^ levels in standard deviations (SD) of baseline. Grayscale shows the Texas Red dextran-labelled vessels. **F)** Astrocytic Ca^2+^ responses occur in proximity to PV_IN_ structures. Colour coding: magenta corresponds to areas of stable high green fluorescence from interneuron GFP; green is dynamic Ca^2+^-dependent fluorescence from astrocytic GCamP6f following optogenetic activation. **G)** The first 10 seconds of the average astrocytic FOV Ca^2+^ responses to optogenetic stimulation in awake mice in (C) calculated in SD of the activity in the entire imaging session. Black lines indicate averages and shaded areas ± SEM. Following optogenetic stimulation of ChR2 of PV_IN_ (red, n=10 FOV, 6 mice), the Ca^2+^ levels were already elevated in astrocytes, opposite to stimulation of VGAT_IN_ (blue, n=8 FOV, 7mice). **H)** The astrocytic Ca^2+^ levels shown in (G) were significantly larger at the termination of the stimulation of PV_IN_ than VGAT_IN_ (t-test, p=0.0091). The dashed line demarcates the threshold of 2 SD to resolve if onset of the Ca^2+^ increase was during stimulation. **I)** The average time to peak for astrocytic Ca^2+^ responses to stimulation was shorter in PV_IN_ than VGAT_IN_ (unpaired t-test, *p<0.05).

We performed the optogenetic experiments in anaesthetized mice and found no change in astrocytic Ca^2+^ (Figure 2C). Next, we stimulated PV_IN_ or VGAT_IN_ in awake mice and detected immediate increases in astrocytic Ca^2+^ (Figure 2C and D; PV_IN_ anaesthetized vs. 2-s stim awake: p=0.0025, t-test, n=10 FOV, 6 mice; VGAT_IN_ anaesthetized vs. 2-s stim awake: p <0.001, t-test, n=8 FOV, 7 mice). These results indicate that the brain state is a decisive factor for interneuron-astrocyte interactions. In awake mice, the automatic active domain analysis detected numerous other changes in astrocytic Ca^2+^, confounding detection of the immediate optogenetic effect (Sup. Figure S2). To further investigate astrocytic Ca^2+^ responses within the FOV, we employed a hypothesis-driven data analysis method that aligns responses to the timing of the stimulation^24^. This allowed us to examine where the Ca^2+^ increases were localized in astrocytes (Figure 2E). Stimulation of PV_IN_ induced an astrocytic Ca^2+^ response (Figure 2E and Sup. Video V1-V3) that primarily aligned with the surface of the interneurons (Figure 2F), whereas VGAT_IN_ stimulation resulted in a sparse and diffuse Ca^2+^ response, optically on the verge of two-photon resolution (Figure 2E). The proximity of the astrocytic Ca^2+^ responses to interneuron structures for increased PV_IN_ activity suggests that the Ca^2+^ responses were due to a direct effect of GABA release from interneurons onto adjacent astrocytic processes. The timing of the astrocytic Ca^2+^ responses to PV_IN_ and VGAT_IN_ stimulation also differed, as the response to PV_IN_ peaked twice and the response to VGAT_IN_ had one peak (Figure 2G). Since Ca^2+^ levels could not be assessed during the 2-second optogenetic stimulation, the onset was deducted from the signal values at the termination of the stimulation. If the first Ca^2+^ level after stimulation was > 2 standard deviations (SDs) of the entire signal, the onset of the response was during stimulation. Accordingly, our data show that the onset of Ca^2+^ responses by stimulation of PV_IN_ occurred during optogenetic stimulation (Figure 2H) and peaked on average 2 seconds after the start of stimulation. In contrast, the astrocytic Ca^2+^ increases induced by VGAT_IN_ occurred over several seconds after termination of stimulation and peaked 4.5 seconds after the start of stimulation (Figure 2I; Student’s unpaired t-test, onset level p=0.0091, peak time p=0.0312; VGAT_IN_ n=8 FOV, 7 mice; PV_IN_ n=10 FOV, 6 mice). The differences in the latency and duration of Ca^2+^ responses evoked by PV_IN_ and VGAT_IN_ suggest that different mechanisms are involved. Optogenetic stimulation of all VGAT expressing interneurons activate PV_IN_ as well as other subtypes of inhibitory neurons. These not only release a mix of peptides besides GABA, but some also inhibit the PV_IN_ ^39^. If the PV_IN_ are inhibited, they will no longer assert their immediate effect on astrocytes, which can explain the loss of the first Ca^2+^ peak in the VGAT_IN_ stimulation.

### Onset of astrocytic Ca^2+^ responses to interneuron activation depends on NA levels

The anaesthetics α-chloralose is an allosteric modulator of the GABA_A_ receptor; therefore, we investigated whether its prohibitive effect on astrocytic Ca^2+^ responses to optogenetic interneuron activation could be mediated by this receptor. To compare we systemically introduced zolpidem (0.5 mg/kg, intraperitoneal [IP], 20 min prior to imaging), which also works as an allosteric modulator of the GABA_A_ receptor^40^ (Figure 3). This sleep-inducing compound was given at a low concentration that did not render the animals to sleep, as assessed by persistent whisker and occasional body movements. Zolpidem had no consistent effect on either the size of the astrocytic Ca^2+^ responses evoked by 2-second stimulation of PV_IN_ or VGAT_IN_ (Figure 3A and B; paired t-test; PV_IN_ p=0.51, n=10 FOV, 6 mice; VGAT_IN_ p=0.36, n=8 FOV, 7 mice), or the onset of the Ca^2+^ responses (Figure 3C and D).

**Figure 3:**
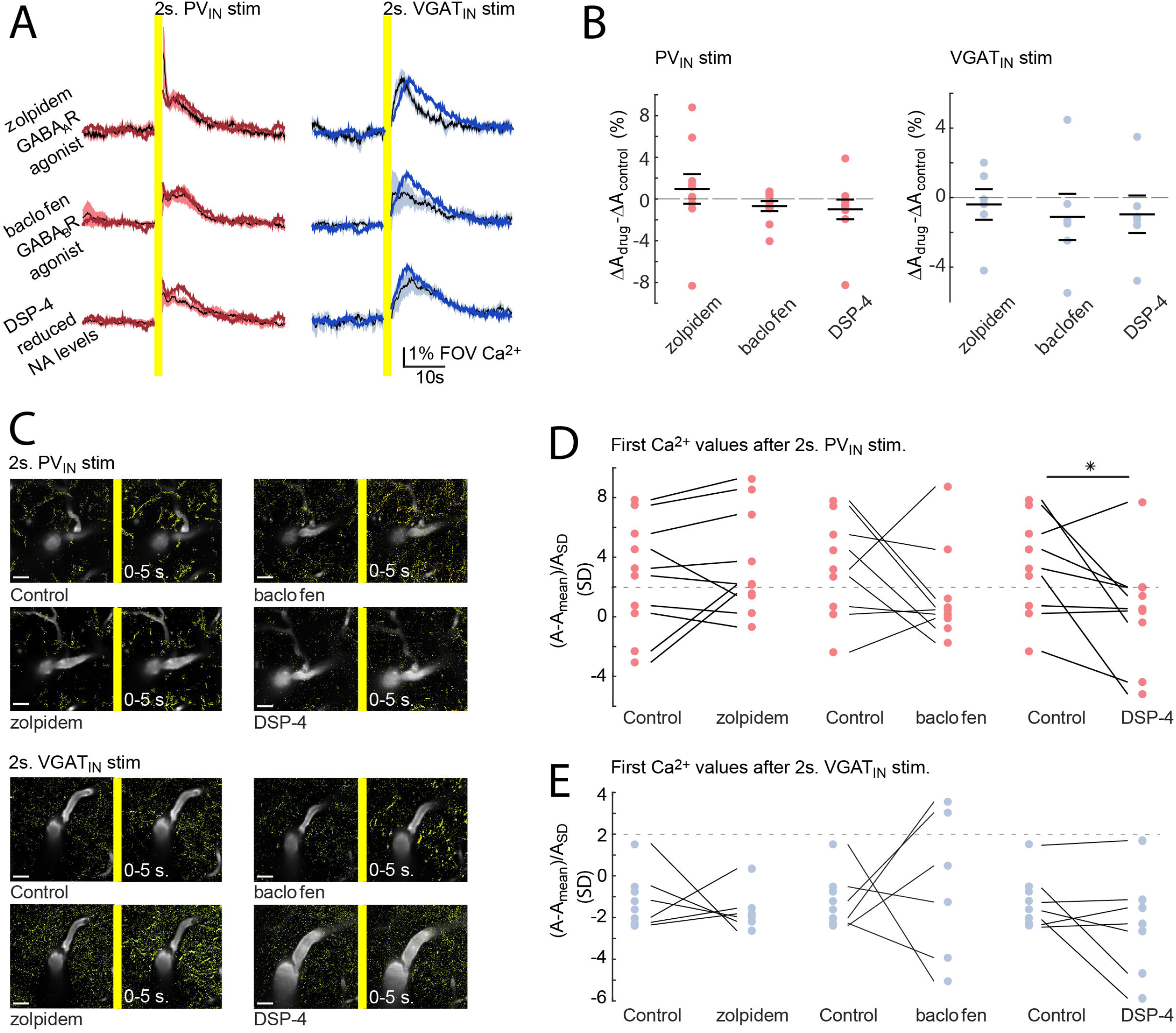
Interneuron-dependent astrocytic Ca^2+^ increases are susceptible to reduced cortical NA levels. **A)** Astrocytic FOV Ca^2+^ responses to 2 seconds of optogenetic stimulation of PV-ChR2-mice (red) and VGAT-ChR2-mice (blue) following GABA_A_R allosteric modulator zolpidem, GABA_B_R agonist baclofen, and NA-reducing DSP-4. Black lines indicate trace averages, shaded areas are SEM, and the colours (red/blue) show the response under control conditions. PV_IN_ n=10 FOV, 6 mice; VGAT_IN_ p=0.34, n=8 FOV, 7 mice. **B)** The peak amplitude of the astrocytic response shown in (A) did not significantly differ after pharmacological intervention. 0.5 mg/kg zolpidem IP: paired t-test, PV_IN_ p=0.51, n=10 FOV, 6 mice; VGAT_IN_ p=0.36, n=8 FOV, 7 mice; 5 mg/kg baclofen: paired t-test, PV_IN_ p=0.18, n=9 FOV, 6 mice; VGAT_IN_ p=0.38, n=8 FOV, 7 mice; 75 mg/kg DSP-4: paired t-test, PV_IN_ p=0.32, n=10 FOV, 6 mice; VGAT_IN_ p=0.34, n=8 FOV, 7 mice. **C)** Ca^2+^ responses to optogenetic stimulation of ChR2 in PV_IN_ (Top) or VGAT_IN_ (Bottom). Pseudocolour coding indicates normalized Ca^2+^ levels in SDs of the baseline for the first 5 seconds after stimulation. The TRITC-dextran labelled vessels are shown in grayscale. Each image pair shows the effect of optogenetic stimulation under control conditions and following drug exposure. Left: Baseline recordings. Right: The respective areas following interneuron stimulation. Scale bars = 20 µm. **D)** Astrocytic Ca^2+^ levels immediately after PV_IN_ stimulation for different pharmacological interventions (zolpidem, baclofen, and DSP-4). The dashed line demarcates the threshold of 2 SD to define whether the onset of the Ca^2+^ increase was during stimulation. The DSP-4 mediated reduction of locus coeruleus-based NA release had a reducing effect (t-test PV_IN_ p=0.038, n=10 FOV, 6 mice). **E)** No effect of either drug on the astrocytic Ca^2+^ levels when VGAT_IN_ stimulation was terminated.

To determine the involvement of the other subtype of GABA receptors, we systemically introduced baclofen, an agonist of the GABA_B_ receptor (5 mg/kg, IP, 20 min prior to imaging). Similar to zolpidem, we observed no effect on the size of the astrocytic Ca^2+^ responses (Figure 3A and B; paired t-test; PV_IN_ p=0.18, n=10 FOV, 6 mice; VGAT_IN_ p=0.38, n=8 FOV, 7 mice), but a trend was found for baclofen reducing the earliest part of the PV_IN_ responses, as only 22% had onset during stimulation (Figure 3C and D).

The activity of the locus coeruleus is the basis of arousal in awake animals^33^. Arousal triggers astrocytic Ca^2+^ increases via brain-wide release of NA^41-43^. Therefore, we excluded recordings if mice ran immediately before, during, or promptly after optogenetic stimulation. In addition, to ensure that the astrocytic Ca^2+^ responses were not due to a startle response related to visual stimulation by the optogenetic fibres, we injected DSP-4 (75 mg/kg, IP, 48 hours prior to imaging), a neurotoxin selective for noradrenergic neurons in the locus coeruleus^32^. Eliminating noradrenergic neurons in the locus coeruleus did not decrease the maximum amplitude of the astrocytic Ca^2+^ responses to optogenetic stimulation (Figure 3A and B; paired t-test; PV_IN_ p=0.32, n=10 FOV, 6 mice; VGAT_IN_ p=0.34, n=8 FOV, 7 mice). On the other hand, DSP-4 treatment affected the onset of astrocytic Ca^2+^ responses to PV_IN_ stimulation. Thus, Ca^2+^ levels at the termination of stimulation were reduced and, judging by Ca^2+^ levels >2 SD, only one response was initiated during the stimulation (t-test PV_IN_ p=0.038, n=10 FOV, 6 mice; Figure 3C and D). This suggests that the early onset of astrocytic Ca^2+^ responses to PV_IN_ stimulation was NA-dependent. Early responses to optogenetic stimulation of VGAT_IN_ were unchanged by DSP-4 (p=0.14, n=8 FOV, 7 mice; Figure 3E). Therefore, it is unlikely that the response to activation of the PV_IN_ is a matter of arousal because the optogenetic stimulation of PV_IN_ and VGAT_IN_ was carried out using the same paradigm. Instead, we propose that part of the astrocytic Ca^2+^ response to activation of PV_IN_ was modulated by cortical NA levels. Thus, reduced NA may partly explain the suppressed astrocytic Ca^2+^ responsivity during anaesthesia and decreased vigilance^31,33,35^.

So far, we had found that the immediate astrocytic Ca^2+^ responses to interneuron activation were faster and more accurate than the Ca^2+^ increases following a GABA puff in anaesthetized mice (Figure 1) and optogenetic interneuron stimulation in brain slices^7^. To elucidate the occurrence of any additional effect of the GABA receptor agonists, we applied our automated active domain analysis to identify long-latency Ca^2+^ responses to interneuron stimulation (Sup. Figure S2). Although we could not detect changes in the long-latency increases in Ca^2+^ under control conditions, following zolpidem treatment, PV_IN_ stimulation increased the number of long-latency astrocytic microdomains (Sup. Figure S2). This finding implies that activation at the GABA_A_ receptor may reinforce the slow fluctuating astrocytic Ca^2+^ responses to GABA.

### Optogenetic stimulation of ChR2 in PVIN and VGATIN induces inhibitory NVC responses in awake mice

To investigate the NVC responses to evoked VGAT_IN_ and PV_IN_ activity, we used intrinsic optical signal (IOS) imaging to assess the extent of the haemodynamic response. We monitored the area of the entire craniotomy (diameter ∼3 mm) (Sup. Figure S3) and found that 2-second optogenetic stimulation of both PV_IN_ alone and VGAT_IN_ induces haemodynamic responses. Vascular responses were widespread across the whole cranial window and were twice as large after activation of VGAT_IN_ than PV_IN_ (Sup. Figure S3B and C; Student’s t-test, p=0.0127, PV_IN_ n=7 mice; VGAT_IN_ n=7 mice).

Next, we reverted to two-photon microscopy imaging to assess the effect of optogenetic stimulation on the diameter changes of PAs and 1^st^ order capillaries adjacent to the astrocytes^44,45^. During anaesthesia, 2-second optogenetic stimulation of VGAT_IN_ induced PA vasodilation, but the effect was absent for stimulation of PV_IN_ alone (Figure 4A). This contrasted with results in awake mice, in which 2-second stimulation of both PV_IN_ and VGAT_IN_ resulted in changes to the PA diameter. Notably, the maximum PA amplitude response was significantly higher for VGAT_IN_ stimulation than for PV_IN_ stimulation (unpaired t-test p=0.0127; PV_IN_ n=10 FOV, 6 mice; VGAT_IN_ n=8 FOV, 7mice). Furthermore, longer stimulation periods significantly increased the PA response amplitude to PV_IN_ but not VGAT_IN_ (paired t-test, PV_IN_ p=0.0383, n=10 FOV, 6 mice; VGAT_IN_ p=0.1788, n=8 FOV, 7 mice; Figure 4B). The PA dilations evoked by 2-second VGAT_IN_ stimulations peaked earlier than with PV_IN_ stimulation (Figure 4C; t-test, p=0.0435). This is in line with previous findings showing that stimulation of VGAT_IN_ triggers the release of vasodilators^46^. Lastly, the vasodilation in response to PV_IN_ stimulation was unaffected by NOS inhibitor L-NNA (Sup. Figure S4). Thus, increases in PV_IN_ activity are likely to mediate dilation through GABA^10^ alone, possibly via a slower and possibly astrocyte-dependent mechanism.

**Figure 4:**
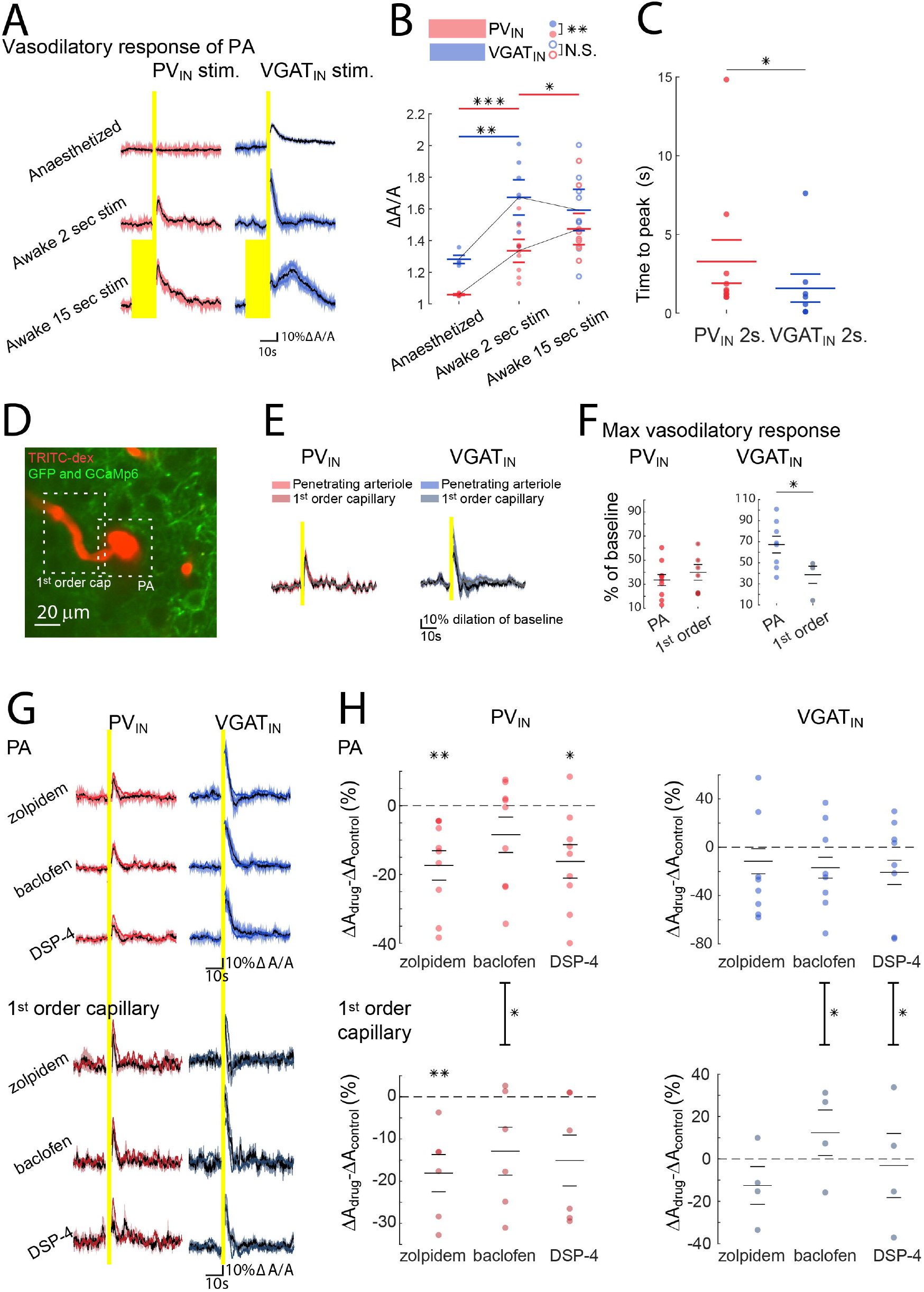
Optogenetic stimulation of awake mice show GABA-dependent NVC. **A)** PA responses to 2-or 15-second optogenetic stimulation of PV_IN_ (red) and VGAT_IN_ (blue) during α-chloralose anaesthesia and in awake mice. Black lines show average traces, shading SEM (PV_IN_ n=10 FOV, 6 mice; VGAT_IN_ n=8 FOV, 7 mice). **B)** VGAT_IN_ 2-seconds stimulations induced larger PA vasodilation than PV_IN_. Response size increased with stimulation of both subtypes of interneurons between anaesthetized and awake state. *p<0.05, **p<0.01, ***p<0.001 (PV_IN_ n=10 FOV, 6 mice; VGAT_IN_ n=8 FOV, 7 mice). **C)** Time to peak comparison between PA responses in PV_IN_ (red) and VGAT_IN_ (blue) stimulation. PA vessels dilated significantly earlier following VGAT_IN_ activation. *p<0.05, unpaired t-test. D) Two-photon image of a PA with branching 1^st^ order capillary. TRITC-dextran in microvasculature (red) ChR2-GFP positive VGAT_IN_ (green) and membrane-bound GFAP-GCaMP6f in astrocytes (green). **E)** Average response of 1^st^ order capillaries (black) and PAs (grey) to PV_IN_ (left) and VGAT_IN_ (right) stimulation (PV_IN_ n=6 FOV, 5 mice; VGAT_IN_ n=4 FOV, 4 mice). SEM shading colour show stimulation and vessel type. (PV=red, VGAT=blue, PA=light, 1^st^ order capillary=dark). F) The 1^st^ order capillaries dilate less than PA to VGAT_IN_ stimulation, whereas dilations are homogenous to PV_IN_ stimulation. *p<0.05, paired t-test (n as in E). **G)** Average PA and 1^st^ order capillary responses to optogenetic stimulation of PV_IN_ (black, SEM:red) and VGAT_IN_ (black, SEM:blue) after pharmacological interventions. From the top: zolpidem (0.5 mg/kg IP), baclofen (5 mg/kg IP), and DSP-4 (75 mg/kg IP) (PV_IN_ n=10 FOV, 6 mice; VGAT_IN_ n=8 FOV, 7 mice). Coloured traces show control response, same as in A). **H)** Changed dilation-size of PAs and 1^st^ order capillaries to PV_IN_ (left, red) or VGAT_IN_ (right, blue) activation. In PV_IN_ stimulation zolpidem and DSP-4 reduced PA dilation and zolpidem reduced capillary dilation. (PV_IN_ PA n=10 FOV, mice 6; capillary n=6, mice 5). None of the drugs had direct effect on dilations in VGAT_IN_ stimulation (PA: n=8 FOV, mice 7; capillary: n=4, mice 4). In PV_IN_ stimulation baclofen reduced 1^st^ order capillary more than PA responses but had the opposite effect in VGAT_IN_ stimulation. Paired t-test *p<0.05, **p<0.01

Next, we focussed on the downstream vascular segment, i.e. the 1^st^ order capillaries branching from the PA (Figure 4D). Vasodilation in response to optogenetic activation of VGAT_IN_ was significantly (∼25%) smaller in capillaries than PAs (paired t-test, p=0.0478, n=4 FOV, 4 mice; Figure 4E and F), whereas the PV_IN_ stimulation equally affected capillaries and PAs (Figure 4E and F). That the activity of VGAT_IN_ did not affect brain microvessels to the same extent may reflect the diversity of processes initiated by such stimulation.

Given that DSP-4 treatment reduced the early rise in astrocytic Ca^2+^, whereas zolpidem and baclofen had no or much smaller effect, we assessed how these drug effects translated to the astrocytic partition in inhibitory NVC. Opposing a singular astrocytic dependence, the allosteric GABA_A_ receptor modulator zolpidem significantly reduced the dilation of both PAs and capillaries following PV_IN_ stimulation (Figure 4G and H; t-test, PA: p=0.0184, n=10 FOV, mice 6; capillary: p=0.009, n=6, mice 5). Zolpidem had no effect on NVC following VGAT_IN_ stimulation, as would be expected from the release of other vasodilators other than GABA (t-test, PA: p=0.2991, n=8 FOV, 7 mice; capillary: p=0.175, n=4, 4 mice; Figure 4G and H). Although others have reported the vasoactive response due to VGAT_IN_ activation in awake mice to have a constriction phase following the dilation^13^, we only observed this constriction after zolpidem administration, but in both PAs and capillaries (Figure 4G). To ascertain the involvement of the GABA_A_ receptor in PV_IN_-dependent NVC, we investigated further using laser speckle and optogenetic stimulation of PV_IN_ in anaesthetized mice. The potential contribution of excitatory synaptic transmission was blocked with local application of the AMPAR and NMDAR inhibitors NBQX (200 µM topically) and MK801 (300 µM topically), whereas the GABA_A_ receptor was targeted by its selective antagonist gabazine (20 µM topically) (Sup. Figure S4). GABA_A_ inhibition was found to enhance NVC, possibly counteracting the effect of α-chloralose. The involvement of the GABA_A_ receptor implies that the curbed NVC response to PV_IN_ stimulation in mice anaesthetized with α-chloralose cannot be explained by reduced astrocytic responsivity alone.

In contrast to zolpidem, the administration of baclofen had no effect on NVC following PV_IN_ or VGAT_IN_ optogenetic stimulation (paired t-test, PV_IN_ PA: p=0.12, n=10 FOV, 6 mice; PV_IN_ capillaries: p=0.072, n=6 FOV, 5 mice; VGAT_IN_ PA: p=0.091, n=8 FOV, 7 mice; VGAT_IN_ capillaries: p=0.29, n=4 FOV, 4 mice; Figure 4G and H). We only found an effect of NA-reducing DSP-4 on NVC for PA dilation following PV_IN_ stimulation; there was no effect with VGAT_IN_ stimulation (paired t-test, PV_IN_ PA: p=0.011, n=9 FOV, 6 mice; PV_IN_ capillaries: p=0.054, n=6 FOV, 5 mice; VGAT_IN_ PA: p=0.081, n=8 FOV, 7 mice; VGAT_IN_ capillaries: p=0.497, n=4 FOV, 4 mice; Figure 4G and H). Importantly, baclofen had a more pronounced effect on reducing the PV_IN_-based dilation in capillaries than PAs (paired t-test, PV_IN_ PA vs. capillaries p=0.011, n=6 FOV, 5 mice; Figure 4H). That the capillary responses were affected only by GABA_B_ receptors is in line with the reduction in astrocytic Ca^2+^ responses observed after baclofen during PV_IN_ stimulation (Figure 3E and D) and in accordance with the notion that astrocytes modulate capillary, but not PA, dilation^29,30^.

In contrast, baclofen had the opposite effect on PAs and capillaries during VGAT_IN_ stimulation, enhancing capillary (paired t-test, VGAT_IN_ PA vs. capillary p=0.017, n=4 FOV, 4 mice) but reducing PA dilatation. Thus, the difference between the PA and capillary response suggests previously unsuspected diverse roles of GABA_B_ receptor in the regulation of NVC at distinct levels of the vascular tree. Lastly, hindering NA release with DSP-4 seemed to affect Pas more than capillaries (paired t-test, VGAT_IN_ PA vs. capillary p=0.021, n=4 FOV, 4 mice). These two preliminary observations may be reflections of some degree of astrocytic contribution to VGAT_IN_ NVC.

### Interneuron-elicited NVC responses correlate with astrocytic Ca^2+^ activity

Although interneuron stimulation increased both astrocytic Ca^2+^ and vessel dilatation in a brain state-dependent manner, our results were ambiguous as to whether astrocytes were required for this inhibitory NVC. The degree of astrocyte involvement was investigated by correlating the Ca^2+^ response to the PA and capillary dilation. In particular, the timing of the astrocytic Ca^2+^ peak relative to VGAT_IN_ stimulation indicates that numerous other components are at play in the inhibitory NVC (Figure 5A), as this lagged behind the NVC responses of the PA by several seconds (paired t-test, p = 0.0094, n=8 FOV, 7 mice; Figure 5B). In comparison, after PV_IN_ stimulation, the astrocytic Ca^2+^ and vasodilation responses did not peak at different times (paired t-test, p=0.3779 n=10, 6 mice). Another indicator of an astrocytic contribution was the linear correlation between the amplitude of the maximal astrocytic Ca^2+^ levels and the size of the PA dilation following PV_IN_ stimulation (Pearson’s R=0.711, p=0.002). Even considering that the amplitude of the delayed astrocytic Ca^2+^ response to VGAT_IN_ stimulation correlated with NVC (Pearson’s R=0.824, p=0.0009; Figure 5C). These correlations were markedly decreased when including the data on pharmacological interventions, which indicated that GABA_A_ receptor activation curbed PA dilation more than it affected the astrocytic Ca^2+^ activity.

**Figure 5:**
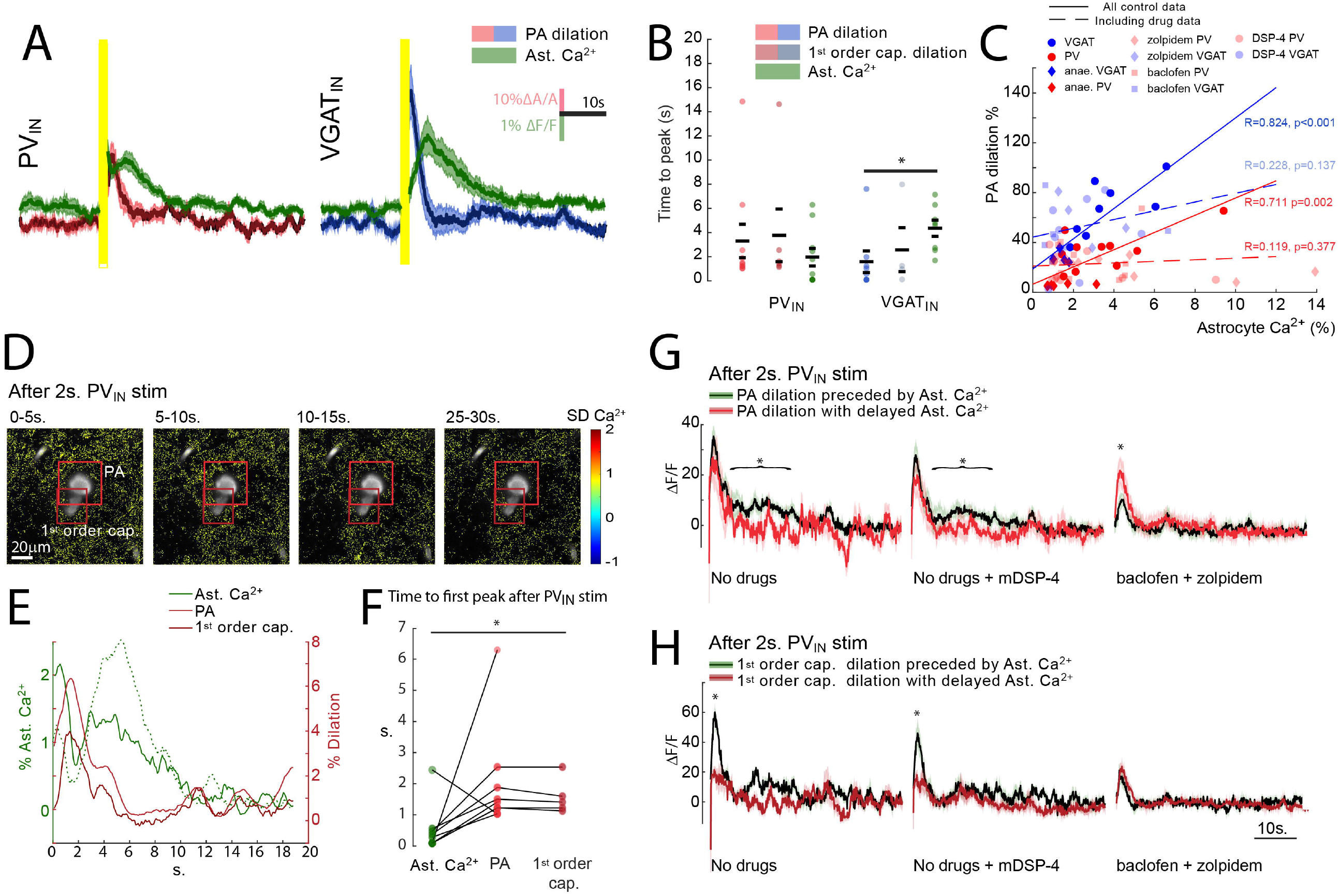
Comparison of NVC and astrocytic Ca^2+^ responses to interneural optogenetic stimulation. **A)** Average PA vasodilatory responses and astrocytic FOV Ca^2+^ responses (SEM: green) following 2-second optogenetic stimulation of PV_IN_ (left, SEM: red, n=10 FOV, 6 mice) and VGAT_IN_ (right, SEM: blue, n=8 FOV, 7 mice). **B)** Time to peak of vasodilatory and astrocytic FOV Ca^2+^ response. In VGAT_IN_ stimulation the astrocytic Ca^2+^ response peaked after vasodilation. *p<0.05. This was not the case in PV_IN_ stimulation. **C)** Correlation of the size of PA dilation and astrocytic Ca^2+^ response. VGAT_IN_ mice, blue. PV_IN_ mice, red. Lines show the linear fit in control conditions, dashed lines the linear fit with inclusion of data from mice after DSP-4, zolpidem, and baclofen (--). Linear correlations were found for both strains under control conditions which were disturbed by pharmacological interventions. D) Example of Ca^2+^ responses to optogenetic ChR2 stimulation in PV_IN_. Ca^2+^ levels are shown in pseudocolours and the vasculature outlined by Texas Red dextran(grayscale). Squares indicate the vessels analysed in (E). E) Changes in astrocytic Ca^2+^ levels and vasculature diameter following PV_IN_ stimulation. F) Timing of the first peak in astrocytic Ca^2+^ levels and of the vessel dilations following PV_IN_ stimulation shows a significant delay between astrocytic Ca^2+^ activity and capillary dilation. p=0.004, paired t-test (n=8 FOV, 6 mice). G) Average PA dilation following PV_IN_ stimulation sorted on the detection of preceding astrocytic Ca^2+^. Black trace from responses preceded by astrocytic Ca^2+^ (threshold: Ca^2+^>2 SD). Red trace from response without (Ca_2+_<2 SD). The significant difference in the second part of the PA dilation for control and +DSP-4 conditions reflect the level of astrocyte contribution. *p<0.05. H) Averaged dilation of 1^st^ order capillaries categorized into two group as in (G). The difference in the first part of the capillary dilation between control experiments and DSP-4 conditions was significantly larger following astrocytic responses. This difference was not preserved following baclofen and zolpidem administration. *p<0.05.

The average trace profiles (Figure 5A) suggested that astrocytic Ca^2+^ transients appear earlier than the NVC response to PV_IN_ stimulation. Analysis of the time to peak of individual responses may reveal either of the two peaks in the Ca^2+^ responses (Figure 5E). Therefore, it was more relevant to focus on the timing of the first peak regardless of its amplitude. In capillaries, the Ca^2+^ response peaked 1-2 seconds earlier than vessel dilation (paired Student’s t-test, PV_IN_ p=0.004, n=5 capillaries, 5 mice; Figure 5F). This supports an early contribution of astrocytic Ca^2+^ in the NVC response to PV_IN_ activation. The onset of the early astrocytic Ca^2+^ response occurred during optogenetic stimulation and can be detected from the astrocytic Ca^2+^ levels exceeding 2 SD upon termination of the stimulation. To identify the significance of the early Ca^2+^ peak for the NVC responses, the PA and 1^st^ order capillary dilations were grouped based on whether it was preceded by this early response in astrocytes (Figure 5G and 5H). We found that the size of the primary dilation of the PA did not depend on astrocytic Ca^2+^ levels, but a later increase was found only in dilations preceded by an early Ca^2+^ peak (t-test, area under the curve (AUC) from 5 to 20 seconds after stimulation; p=0.0373, n=10 FOV, 6 mice; Figure 5G). The same pattern was found when including dilation in the DSP-4-treated animals (t-test, AUC 5-20 seconds after stimulation; p=0.0381, n=10 FOV, 6 mice). Following zolpidem or baclofen, the first part of the PA dilation was reduced in size, but this reduction occurred in responses preceded by astrocyte activity. Modulated by either the GABA_A_ or GABA_B_ receptor agonists, the dilations occurring after astrocytic Ca^2+^ peaks were much reduced in size and significantly smaller than those occurring without early astrocytic Ca^2+^ (t-test, max level from 1-5 s after stimulation p=0.025, n=10 FOV, 6 mice; Figure 5G). A similar investigation in the dilation of 1^st^ order capillaries in response to PV_IN_ stimulation revealed a stronger dependence on astrocytic Ca^2+^ timing (Figure 5H). At this part of the vasculature, the primary part of the dilation was much larger when preceded by an early astrocytic Ca^2+^ peak (t-test, max levels from 1-5 s after stimulation p=0.019, n=10 FOV, 6 mice; Figure 5G) and similarly when including the data from the mice with DSP-4-reduced cortical NA levels (t-test, max levels from 1-5 s after stimulation p=0.026, n=10 FOV, 6 mice). These correlations imply a strong contribution of this early astrocytic Ca^2+^ increase in the dilation of 1^st^ order capillaries. The reduced capillary dilation following zolpidem and baclofen treatment appeared to affect the responses preceded by early astrocytic Ca^2+^. This effect suggests that the dilations reduced by these drugs are those supported by astrocyte activity. With the introduction of GABA_A_ or GABA_B_ receptor agonists, the astrocyte-based dilation of capillaries was reduced to the level observed without early astrocytic Ca^2+^ activity (Figure 5H). These results indicate that the basic capillary dilation in response to PV_IN_ stimulation can be increased by astrocytic Ca^2+^ responses, but not following zolpidem or baclofen exposure. We conclude that astrocytic Ca^2+^ signalling augments PV_IN_ -dependent NVC responses, especially in capillaries.

## Discussion

Astrocytic involvement in neural circuits has recently been recognized but, thus far, most studies have focused on the interaction of astrocytes in the context of glutamatergic signal transmission^47^. As inhibitory interneurons regulate excitatory activity and shape network function, it is important to elucidate whether the roles of astrocytes in neuronal circuits also include direct and indirect modulation of inhibitory processes. Here, we showed that astrocyte responsivity to GABA depends on the awake state of the animal, which is in line with previous observations of brain state dependence^31,34,35^. Using awake mice, we found that astrocyte activity may link interneuronal GABAergic signalling to NVC, particularly the vascular responses of capillaries.

Astrocytic Ca^2+^ responses evoked by increases in PV_IN_ activity occurred earlier than responses evoked by VGAT_IN_. The delayed response to the VGAT_IN_ stimulation compared to PV_IN_ could reflect the activation of a mixed population of interneurons including the interneuron subtype VIP, which is known to inhibit PV_IN_^39^. Another explanation could be a combination effect, as stimulation of VGAT_IN_ will release GABA and several neuropeptides, in contrast to PV_IN_ releasing only GABA. These two stimulation paradigms reflect the complex interneuron-astrocyte interactions, which may explain the diversity of astrocytic Ca^2+^ responses reported in the literature^5,8,48^. The GABAergic astrocytic Ca^2+^ responses were immediate and transient but could spontaneously fluctuate under non-physiological conditions, such as in response to local puffing of GABA or after modulating the GABA_A_ receptor activity. Such stimulation could be considered supra-normal^49^ compared to our brief optogenetic responses. Long-lasting increases in delayed, spontaneously fluctuating astrocytic Ca^2+^ have been described to depend on both GABA_A_ and GABA_B_ receptors and neuropeptides in several slice studies and in one in vivo study using anaesthetized mice^5-8,50^. Astrocytic Ca^2+^ responses to GABA_B_ receptor binding are IP3-dependent^48,51^, but no mechanisms have yet been suggested for potential subtype-specific, neuropeptide-dependent effects. Our study is the first to show that the long-lasting effect of GABA was curbed by the neuropeptide CCK. Cortical pyramidal neurons^52^ and astrocytes express CCK receptor, as a few studies have reported a direct effect of this neuropeptide^53,54^. We found that CCK reduced the GABA effect on astrocytes, where another neuropeptide, somatostatin, has been found to potentiate a GABA-induced response^7^. These observations suggest a complex relationship between GABA-mediated astrocytic Ca^2+^ activity and interneuron subtype-specific responses.

In the cortex of awake mice, GABA from PV_IN_ triggered an immediate Ca^2+^ increase in astrocytes. Neither the GABA_B_ receptor agonist baclofen nor the allosteric GABA_A_ receptor modulator zolpidem significantly modified the immediate interneuron-dependent astrocytic responses. In slices and anaesthetized animals, baclofen potentiates astrocytic GABA_B_ -based IP_3_ -dependent Ca^2+^ increases^8,48^, but in awake animals we found that baclofen more likely delayed the Ca^2+^ response in astrocytes. Therefore, it is also possible that the delay in astrocytic Ca^2+^ responses to VGAT_IN_ activation is due to more GABA_B_ receptor activation than with PV_IN_ activation alone. Notably, the effect of zolpidem on Ca^2+^ increases following PV_IN_ activation appears to be bimodal. Occasionally, astrocytic Ca^2+^ responses were enhanced as would be expected from the combined action of PV_IN_-derived GABA release and the allosteric modulator on GABA_A_ receptor. However, most Ca^2+^ responses were decreased, similar to the effect of α-chloralose, which also exerts its actions as a allosteric modulator of the GABA_A_ receptor^55^. Though our use of zolpidem did not render the animals to sleep, the animals did appear to be less active and the variability in Ca^2+^ responses after zolpidem may depend on the activity level of each individual animal. The astrocytic Ca^2+^ response to GABA may be due to GABA receptor binding and/or astrocytic GABA uptake by the GABA transporter GAT3^51,56^, which could then modify intracellular Ca^2+^ through the reverse mode of the Na-Ca^2+^ exchanger^51,57^. We found that neither the GABA_A_ nor GABA_B_ receptor agonists significantly modified astrocytic Ca^2+^ levels, supporting that the GABA transporter contributes to the effect. Abolition of NA release reduced astrocytic Ca^2+^ responses to PV_IN_ activation. This supports the lack of astrocytic responsivity to interneuron activation in the anaesthetized animals being due to the brain state change and reduced cortical NA levels^58^. Similarly, it is plausible that the weak effects we observed after targeting the GABA_A_ and GABA_B_ receptor may be primarily systemic rather than directly on astrocytes.

The immediate Ca^2+^ increase in response to interneuron activation in awake mice was localized to small domains compared to arousal-based activity. Thus, it was not well detected, even with the optimized event detection algorithm based on changes in single-pixel intensities. Our automated microdomain detection method was similar to analytical techniques developed in other studies to quantify astrocyte activity^35,38^. Nevertheless, the response to the optogenetic stimulation was evident. The perisynaptic processes of astrocytes are very fine and well below the diffraction limit of traditional fluorescence microscopy^2,59^. Therefore, smaller Ca^2+^ responses may remain undetected. We found that analysis of astrocytic responses was more precise using a post-hoc method established in our previous study^24^, and that it localized the astrocytic responses to areas along the interneurons expressing ChR2. This highlights the importance of the choice of analytical methods in studies concerning astrocytic Ca^2+^ activity.

Inhibitory NVC was also severely altered in awake mice compared to anaesthetized animals. Under anaesthesia, we found that activation of VGAT_IN_ triggered a NVC response that was greater in awake mice. Furthermore, VGAT_IN_ evoked an NVC response that peaked during or just after stimulation. Among VGAT_IN_, several express nNOS and are expected to release the vasodilator NO upon activation^10,11^, which can account for the first part of the CBF response to interneuron activation^46^. VGAT_IN_ activation releases other vasodilators, as well as vasoconstrictors, such as VIP or NPY^12^. Based on the reduced dilation under anaesthesia, the release patterns of these vasoactive compounds must be brain state-dependent. Though we did see a correlation between astrocytic Ca^2+^ responses and VGAT_IN_ NVC, the many pathways recruited in this optogenetic stimulation makes it difficult to hypothesize astrocytic involvement. We hypothesize that NVC responses triggered by VGAT_IN_ stimulation reflect the combined release of GABA, vasoactive factors, and a mixture of additional neuropeptides.

We found that our anaesthetics blocked the PV_IN_ NVC in 2-second optogenetic stimulation. In contrast, in awake mice, we persistently observed a vascular response to PV_IN_ stimulation, which is in contrast to one report of PV_IN_-based cerebral vasoconstriction^60^ but in line with previous studies^14^ that explained the disparities as a difference in stimulation paradigms. We also observed that PV_IN_ triggered an NVC response that increased after stimulation and peaked several seconds later. PV_IN_ primarily release GABA upon activation and contain only small amounts of NOS^10^. As PV_IN_ stimulation-based NVC was insensitive to NOS inhibitors, we assume that the CBF responses were triggered by GABA release^15,19^. It is unlikely that GABA targets receptors on vascular mural cells (i.e., vascular smooth muscle cells and pericytes) because the gene expression of GABA_A_ receptor and GABA transporter genes^61^ in pericytes is weak and 100-1000 orders of magnitude lower than in astrocytes. The difference in NVC responses to interneuron activation between anaesthetized and awake mice points to brain state dependence. Similar to the repressed astrocyte activity^31^, PV_IN_ reduce their spiking activity during down-states^62^. It is possible that the reduction in spikes is reflected in reduced GABA release. This reduction cannot be absolute since we found intact NVC in anaesthetized animals when using longer optogenetic stimulation^15^. In addition, any reduced GABA release would be inconsequential because we found that astrocytic sensitivity to GABA was reduced in the anaesthetized animals with direct puffing. In awake mice, astrocytes were very responsive to PV_IN_ stimulation in a manner that preceded the vascular response and correlated well with capillary dilation. As astrocytes express GABA receptors and transporters in addition to secreting vasoactive compounds^20-22^, we hypothesize that the interaction of GABA with astrocytes facilitates PV_IN_-evoked NVC via astrocytic release of vasodilators.

The timing of astrocytic Ca^2+^ and CBF and the immediacy of responses suggest that astrocytes do not contribute substantially to the VGAT_IN_ NVC responses. In comparison, GABA release is likely a major factor in PV_IN_ NVC responses and vasodilation possibly aided by astrocytic Ca^2+^. Further studies are needed to pinpoint a specific mechanism, but our results suggest that astrocytic effects are sensitive to segmental differences in the microvasculature, where its effects are more strongly exerted on capillaries than on PAs^29,30^. In conclusion, we propose that astrocytes react rapidly to GABA and may contribute to PV_IN_ - dependent NVC. Our results emphasize the interplay between astrocytes and inhibitory neurons, which depend on both GABA and neuropeptides, and reveal that astrocytes may take part in the control of NVC evoked by specific GABAergic interneurons.

## Methods

### Animals

All procedures were performed in compliance with the guidelines of the European Council’s Convention for the Protection of Vertebrate Animals Used for Experimental and Other Scientific Purposes and approved by the Danish National Ethics Committee. All experiments were performed on WT C57BL/6J, VGAT ChR2-EYFP, and PV-ChR2-EYFP mice on a C57BL/6J background. Animals were at least 8 weeks old at the time of chronic window implantation and viral vector injection, and all experiments were performed on animals aged between 10 and 38 weeks. All animals were injected with 3×200 nL AAV5-gfaABC1D-lck-GCaMP6f^37^. Both male and female animals were used. Animals were group-housed both prior to and after chronic window implantation with access to food and water ad libitum on a 12/12-hour night/day cycle.

### Surgical procedures for acute experiments

Surgical procedures for acute experiments were carried out as described previously^24^. Anaesthesia was initiated using IP ketamine/xylazine (xylazine 10 mg/kg; ketamine 60 mg/kg, then 30 mg/kg every 20–25 min). Animals were maintained at 37 □ on a heating pad with a rectal thermometer probe. First, they underwent tracheotomy for mechanical respiration (180–220 µL volume; 190–240 strokes/min) with O_2_-supplemented air (1.5–2 mL/min, MiniVent Type 845 ventilator, Harvard Apparatus). The left femoral vein and artery were then cannulated. Intravenous (IV) access was used for anaesthetic and vascular dye delivery. After life support surgery, the scalp was removed and the skull glued (Loctite Adhesives) to a custom-made metal plate. A craniotomy was drilled (3 mm lateral, 0.5 mm posterior to bregma, Ø=3 mm) and the dura carefully removed before covering the brain surface with 1% agarose (type III-A; Sigma-Aldrich) and artificial CSF (in mM: KCl 2.8, NaCl 120, Na_2_HPO_4_ 1, MgCl_2_ 0.876, NaHCO_3_ 22, CaCl_2_ 1.45, glucose 2.55; at 37□, aerated with 95% air/5% CO_2_ to pH 7.4). A glass coverslip (4×4 mm, #1.5 thickness; Thermo Scientific) was positioned over two-thirds of the craniotomy, leaving a small gap for electrode insertion. After surgery, the anaesthesia was switched to α-chloralose (50 mg/kg/h IV) and the animal allowed to rest for 25 min before imaging. Throughout imaging, we monitored end-tidal respiratory CO_2_ levels (Type 340 capnograph; Harvard Apparatus) and arterial blood pressure via a femoral artery catheter (BP-1 Monitor, World Precision Instruments). Prior to imaging, an arterial blood sample (∼55 μL) was collected for blood gas evaluation (ABL 700 Series, Radiometer) and, if necessary, the respiration was adjusted.

### Viral vector injections

Viral vector injections were performed following our adaptation of standard procedures^25^. The procedure was performed under aseptic conditions and isoflurane anaesthesia (5% induction, 1.5% during surgery in 10% O_2_ in air mixture). The fur on top of the head was shaved off and the animal placed in a stereotaxic frame on a heating pad to maintain a body temperature of 37□. The eyes were kept moist using eye ointment (Viscotears, Novartis). The shaved skin was disinfected with chlorhexidine/alcohol (0.5%/74%; Kruuse). Carprofen (5 mg/kg body weight [BW]; Norodyl, Norbrook) was administered subcutaneously, and 0.5 mL saline was distributed over several sites to keep the mouse hydrated during surgery. Lidocaine (100 µL 0.5%) was subcutaneously injected under the scalp. A cut was made in the skin over the right side of the skull and the periosteum removed using sterile cotton swabs. A small burr hole was drilled within the intended site for the craniotomy. Three doses (200 nL) of AAV were deposited using a glass micropipette. After injection, the pipette was slowly retracted, the hole in the skull closed with bone wax (Surgical Specialties), and the skin glued together again using GLUture topical tissue adhesive (Zoetis). Next, the animal was returned to the cage on a pre-warmed heating pad to wake from anaesthesia. The cage contained pre-wetted food pellets for easy eating and hydration. Postoperative care consisted of subcutaneous injections of Rimadyl (24 and 48 h post-surgery, dose as before). The convalescent animals were carefully monitored for 7 days post-surgery. Mice were injected with a commercial AAV construct to express the Ca^2+^ indicator GCaMP6f in astrocytes under control of the gfaABC1D promotor. An insert was added to the construct such that the Ca^2+^ indicator was tethered to the cell membrane to better assess the activity in small perisynaptic processes^37^.

### Chronic window implantation

Chronic window implantation was performed under aseptic conditions and isoflurane anaesthesia (5% induction, 1.5% during surgery in 10% O_2_ in air mixture). The fur on top of the head was shaved and the mouse placed in a stereotaxic frame on a heating pad to maintain a body temperature of 37□. The eyes were kept moist using eye ointment (Viscotears, Novartis). The shaved skin was disinfected with chlorhexidine/alcohol (0.5%/74%; Kruuse). Carprofen (5 mg/kg BW; Norodyl, Norbrook) and buprenorphine (0.05 mg/kg BW; 19 Temgesic, Indivior) were administered subcutaneously, and 0.5 mL saline was distributed over several sites to keep the mouse hydrated during surgery. Lidocaine (100 µL 0.5%) was subcutaneously injected under the scalp. The scalp was removed and the surface of the skull cleaned and the periosteum scraped away using a scalpel. Craniotomy was performed over the right somatosensory cortex (3 mm lateral, 0.5 mm posterior to bregma; Ø=3 mm). The bone flap was carefully removed and the exposed brain temporarily covered with a haemostatic absorbable gelatine sponge (Spongostan®, Ferrosan, Denmark) pre-wetted with ice-chilled aCSF. The cranial opening was filled with aCSF, and then sealed with an autoclave sterilized round imaging coverslip (Ø=4 mm, #1.5 thickness; Laser Micromachining LTD). The rim of the coverslip was secured with glue and dental cement, and a lightweight stainless steel head plate (Neurotar®) was positioned on the top of the skull surrounding the cranial window. The skull was coated with glue, followed by adhesive resin cement (RelyX Ultimate, 3M) to secure the exposed bone, including the skin incision rim, and firmly attach the metal plate to the head. After the surgery, the animals were injected with dexamethasone (4.8 mg/g BW; Dexavit, Vital Pharma Nordic) and returned to the cage on a pre-warmed heating pad to wake from anaesthesia. They were provided with pre-wetted food pellets for easy eating and hydration. Postoperative care consisted of subcutaneous injections of carprofen and buprenorphine (24 and 48 h post-surgery, dose as before). The convalescent animals were carefully monitored 7 days post-surgery. When fully recovered from the surgery, habituation training began. The animals were gradually habituated to the experimenter handling, the mobile home cage (Neurotar®), head fixation, and imaging. Sugar-sweetened water was used as a reward during training.

### Pharmacology

For imaging experiments in awake mice, drugs were delivered as a single IP injection approximately 15 minutes before imaging commenced (baclofen and zolpidem), or >48 h before imaging commenced in the case of DSP-4. R-baclofen (Tocris) was dissolved in saline and a single dose of 5 mg/kg BW administered. Zolpidem was dissolved in a 10% DMSO/saline solution and a single dose (0.5 mg/kg BW) administered. DSP-4 (Tocris) was dissolved in saline and a single dose (75 mg/kg BW) administered.

For acute imaging experiments in anaesthetized mice, drugs were delivered locally via intracortical injection using a glass micropipette as described previously^36^. GABA was dissolved in aCSF (in mM: NaCl 120, KCl 2.8, Na_2_HPO_4_ 1, MgCl_2_ 0.876, NaHCO_3_ 22, CaCl_2_ 1.45, glucose 2.55, at 37°C, pH 7.4) at a concentration of 50 or 500 μM. To visualize the injection during two-photon acquisition mode, the injectant was mixed with Texas Red in a final dye concentration of 100 μM. In a few experiments, the following drugs were added topically to the craniotomy 20 min prior to optogenetic stimulation: NOS inhibitor L-NNA (1 mM), GABA_A_ receptor agonist gabazine (20 µM), NMDAR antagonist MK801 (300 µM), and AMPAR antagonist NBQX (200 µM).

### Two-photon imaging

All two-photon imaging experiments were carried out on a Leica TCS SP5 MP, DM6000 CFS system equipped with a MaiTai Ti:Sapphire laser (Millennia Pro, Spectra-Physics) using a 20X 1.0 NA water-immersion objective. The emitted light was split by a dichromatic filter at 560 nm into two channels and collected by separate multi-alkali detectors after 500–550 nm and 565–605 nm bandpass filtering (Leica Microsystems). Animals were injected through a venous catheter (acute experiments) or retro-orbitally under isoflurane anaesthesia (awake experiments) with a 2.5% weight/volume solution of Texas Red dextran (70 kDa) in saline. Imaging laser power was kept below 35 mW at all times to prevent induction of astrocyte activity due to phototoxicity. Images were acquired from cortical layers I-III.

### Optogenetic stimulation

Optogenetic stimulation at a wavelength of 473 nm was delivered from a diode-pumped solid-state laser (Crystalaser, DL473-100-O) through an optical fibre (Thorlabs; 0.22NA Ø multimode, ceramic ferrule 1.25 mm) to the brain. The fibre was guided from the side between the cranial window and the objective. The fibre was positioned at an approximately 30 degree angle to the plane of the cranial window at a distance of ∼4 mm, resulting in an elliptical illuminated area with long and short axes of approximately 2.125 mm and 0.9 mm that included most of the cranial window. The stimulation power was 5 mW at the cranial window. The stimulation was a pulse train of 2-or 15-second total duration at a frequency of 100 Hz and duty cycle of 75% corresponding to a pulse duration of 7.5 ms.

### Data analysis

All analyses were performed using Matlab R2017b. In two-photon imaging, all fluorescence recordings were motion-corrected before further analysis.

The Neurotar mobile HomeCage® allows for locomotion tracking. The movement speed of displacement recorded by the Neurotar mobile HomeCage® was thresholded at 1 cm/s for binary movement information (locomotion/no locomotion).

Intrinsic Optical Signal (IOS) videos were analysed using ImageJ/Fiji. Frames up until stimulation were averaged to yield a baseline image, and all frames were then divided pixel by pixel by the baseline image. The average intensity over the whole craniotomy was calculated to extract single time series traces.

Vasodilation was calculated as dA/A. Penetrating arterioles and 1^st^ order capillaries were manually selected and a threshold selected. Fluorescence videos of blood vessels were spatially smoothed using a 3×3 pixel median filter that preserves sharp edges while denoising better than a Gaussian filter. Masks for vessels of interest were drawn by hand. All frames were filtered by the triangle thresholding method. The now binary images were cleaned up using morphological opening and closing, and the foreground area was calculated. For normalized dilation traces, the median of the whole trace was used as a baseline. The area was calculated and normalized by the median of the area trace for a whole recording.

Ca^2+^ microdomain detection focused on changes in fluorescence rather than the strength of fluorescence. The method consists of thresholding followed by the clean-up of active voxels (two spatial dimensions and one temporal dimension), and then clustering of detected active voxels and analysis of these clusters (Sup. Figure 1). Although there are key differences, the detection scheme outlined below is not too dissimilar to AQuA^38^. The algorithm identified changes in individual pixel intensities by noise-based thresholding. In this way, we circumvented challenges related to differences in background fluorescence intensities that may confound the data analysis^63,64^. Thresholding was based on two parameters: the baseline (µ) estimated with the mode and the baseline variation (σ). A mode of the raw trace underestimates the baseline level because many pixels would often show no signal and, therefore, lie at the minimal fluorescence intensity value possible. Instead, we used the value around which the signal fluctuates at rest. In this step, we were only interested in the level around which the signal fluctuates and not the fluctuations themselves so we could use the smoothed data. Therefore, signals from each pixel were low pass-filtered using a 5^th^ order Butterworth filter with a cut-off frequency of 0.26 Hz (5% of the Nyquist rate) (Sup. Figure S1B). The data were binned to a 50-count width; the fluorescence signals are on the order of 10,000. This binning sacrifices some resolution to circumvent the issue of the minimal value appearing several times. By binning the signal, the value around which the signal fluctuates should be better represented in relation to the single minimal value around which the signal does not fluctuate. After filtering and binning, the mode was found as an estimate of the baseline. An estimate of the baseline variation around the baseline level was found by high pass filtering the signals again using a Butterworth filter with the same cut-off frequency as for the low pass filter. By doing this, we discarded the larger changes stemming from actual Ca^2+^ activity within the cell and obtained only the high frequency noise through which the signals need to stand out. As a first rough estimate of the baseline variation, we took the SD of this high pass-filtered trace. Finding the variance by using the SD of this signal results in an overestimation because the noise in the fluorescence is due, in part, to Poisson processes in both the fluorophores and the detectors. For the Poisson distribution, the variance is equal to the mean (µ=σ2); thus, with greater signal strength, we obtain greater variation. For this reason, we used a variance stabilization strategy based on iterations. To find the baseline, we threshold the signal with our current estimates (the baseline level found as described above and the baseline variation represented by the SD of the high pass-filtered signal). Anything below a threshold of µi + 2 · σi (the i’th pixel) was considered a true baseline. We then used this new trace, from which we have discarded a large portion of the Ca^2+^ signals, to better estimate the baseline level and baseline variation (Sup. Figure S1D). This estimation of baseline level and variation was used to threshold each pixel signal (Sup. Figure S1E). Next was a clean-up of the detected active voxels. Even after a threshold of two sigma, there will always still be some false-positives; thus, lone voxels without any active neighbours were disregarded, and voxels that were close together were grouped. As the data are a binary 3D set (spatiotemporal, Sup. Figure S1F), grouping was accomplished by using simple morphological operations commonly used for noise removal in computer vision: opening and closing (Sup. Figure 1G).

In the next step, the active voxels were clustered into domains that are active together. Clustering was achieved in 2D but, to avoid clustering events that might partially or wholly be within the same spatial region but do not overlap in time, the binary 3D array was projected into 2D first. Thus, the overlapping clusters were projected onto the same or connected pixels and consequently clustered together. This may result in an underestimation of the amplitude of spatially small signals that happen to lie within a larger cluster because the average fluorescence of the whole area is calculated for each frame and, if only part of the cluster is active in some frames, the signal will be underestimated.

These metrics yield the total area of the FOV that was active during the recording, the total integrated fluorescence of all signals within the recording, and the number of spatial regions where activity was present. The ΔF/F traces from these clusters can then be further worked on (Sup. Figure S1I). The accuracy of this detection algorithm was tested on synthetic data produced at various levels of contrast and signal-to-noise ratio (SNR).

For the hypothesis-based analysis, we combined recordings from individual optogenetic stimulations into one meta-recording. The resulting meta-recording arose from reorganizing the images in time relative to the onset of the optogenetic stimulations. The images were then normalized over time to obtain the pixel-based difference in fluorescence levels independent of the level of stable GFP expression in the interneurons. For quantitative and illustrative purposes, the normalized fluorescence changes were then averaged in 5-second intervals following the onset of optogenetic stimulation. The areas with stable high fluorescence in the green channel were identified as interneuron structures (magenta in Figure 2G), and the areas with the strongest change in fluorescence following optogenetic stimulation were considered to be the astrocytic Ca^2+^ responses (green in Figure 2G).

### Statistical analysis

Normality was tested visually and data log-transformed when appropriate. Mixed-effects models were used to test significance, followed by two-tailed paired or unpaired Student’s t-test as appropriate with Holm-Sidak correction for multiple comparisons and Welch’s correction in cases of unbalanced groups. Significance was set at 0.05.

## Supporting information

Four supplemental figure

## Acknowledgements

We would like to acknowledge our animal technician Micael Lønstrup for his help with animal experiments. We also thank Anna Devor at UCSD and her laboratory for training optogenetics in awake mice and Krzysztof Kucharz for his useful input on the manuscript. This study was supported by “Læge Sofus Carl Emil Friis og Hustru Olga Doris Friis’ Legat”, “Dagmar Marshalls Fond”, and “Hørslev fonden”.

## Author’s contribution

A.K. and B.L. designed the study. A.K., L.S., and M.D. performed the experiments. A.K., G.V., K.T., and B.L. analysed the data. A.K. and B.L. wrote the paper with input from M.L. and all authors.

## Data availability

The data supporting these findings are available from the corresponding author upon request.

## Code availability

Detection scripts are available online on GitHub, and all other code is available from the corresponding author upon request.

---

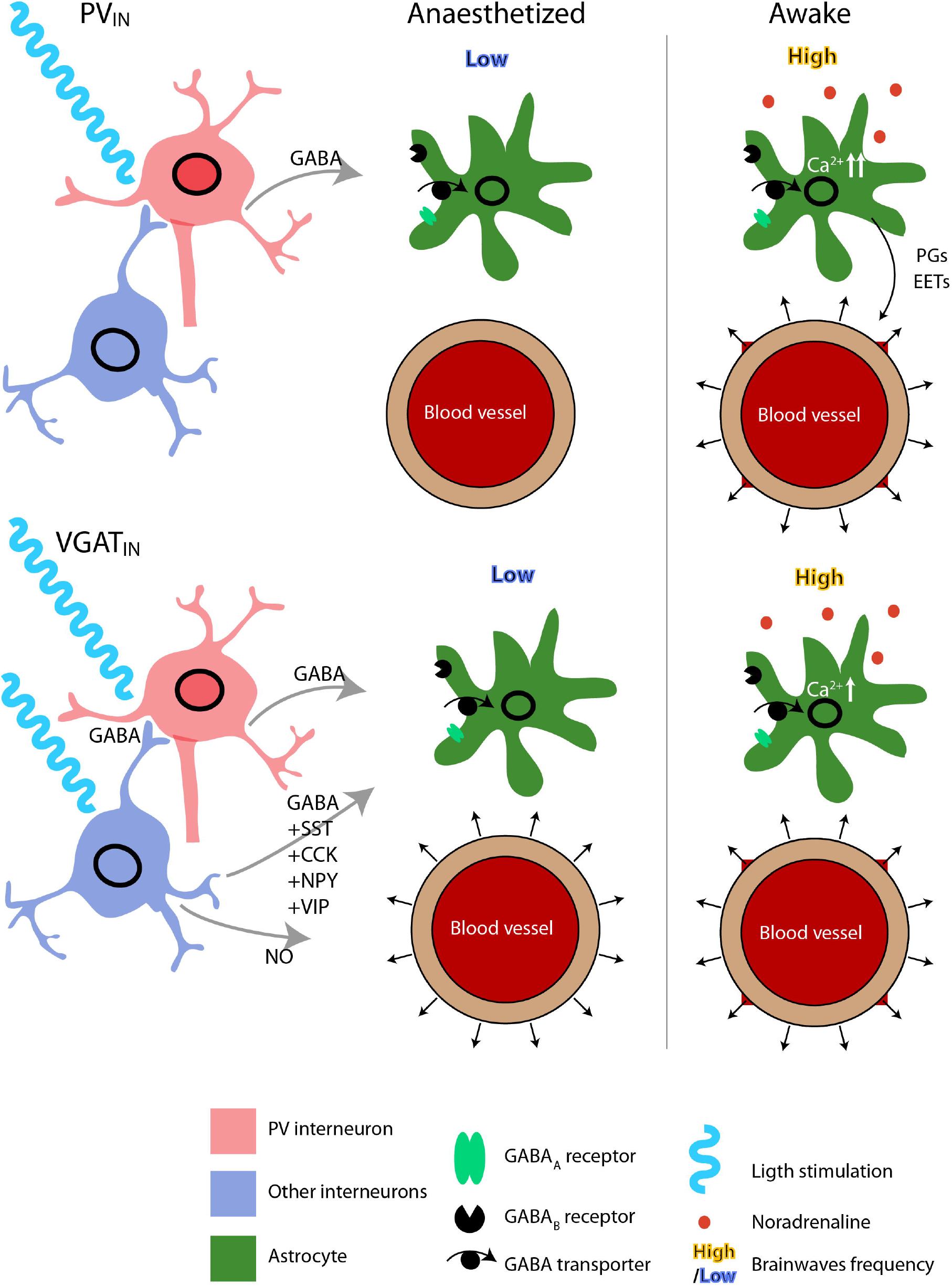

