## Supplementary material for "Astrocytic Ca^2+^ signals partake in inhibitory neurovascular coupling in a brain state-dependent manner": Four supplemental figure

**Supplemental information for** Astrocytic $Ca^{2+}$ signals partake in inhibitory neurovascular coupling in a brain state-dependent manner.

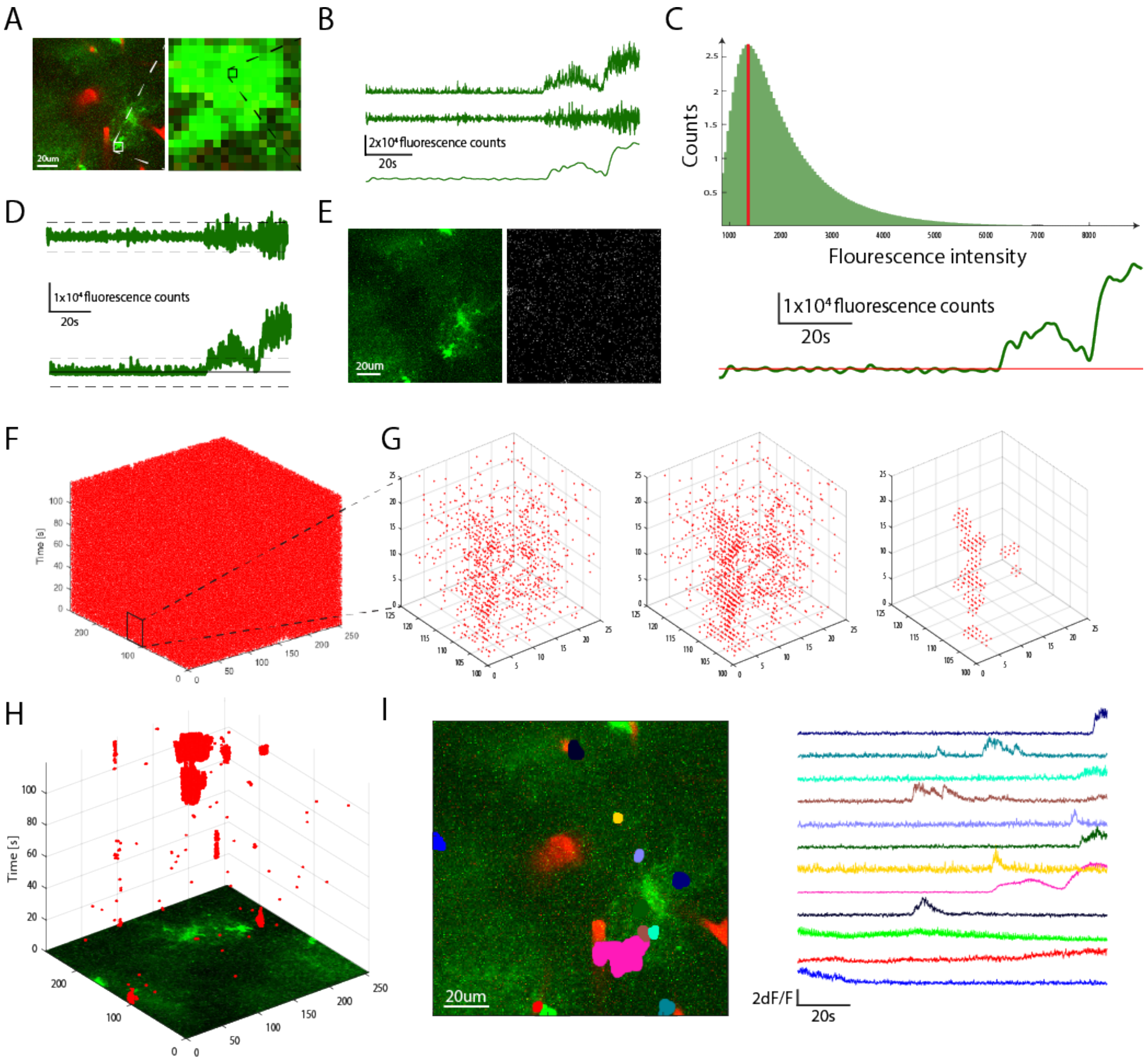


**Supplementary Figure S1:**

**Microdomain $Ca^{2+}$ event detection algorithm.**

**A)** Example frame from a two-photon recording. Each pixel is analysed for changes in signal intensity. The detected active voxels are later combined into domains and analysed together.

**B)** **Top:** Raw signal from the pixel highlighted in (**A)**. **Middle:** Signal after high pass filtering. **Bottom:** Signal after low pass filtering.

**C)** **Top:** Histogram of signal values from the low pass filtered signal. The mode is highlighted in red. **Bottom:** Mode plotted on top of the low pass filtered signal. The mode is used as an estimate of the baseline level.

**D)** **Top:** High pass filtered signal is used to estimate the baseline variation in SD. **Bottom:** Baseline level and baseline variation estimates plotted on top of the raw signal.

E) All frames (example frame on the left) are thresholded using estimates of the baseline level and variation resulting in binary images (right).

**F)** Each recording then becomes a binary 3D (xyt) array. The thresholding will inevitably result in some false positives due to noise.

**G)** **Left:** Zoom-in on a small subvolume of the whole recording. **Middle:** The same subvolume after clean-up using morphological closing. **Right:** The same subvolume after clean-up using morphological closing first, and then morphological opening.

**H)** Resulting binary 3D array showing detected events. Active voxels are clustered by connected components into active domains.

**I)** **Left:** Active astrocytic domains detected within a FOV. **Right:** Corresponding normalized fluorescence traces from the detected domains.

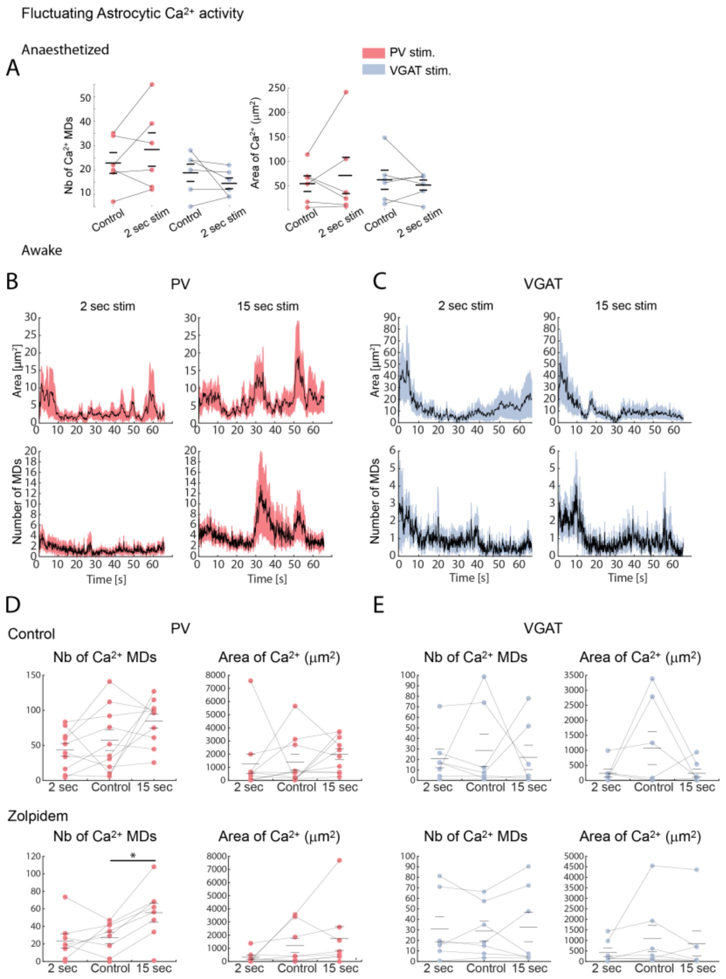


**Supplementary Figure S2:**

**Microdomain analysis of astrocytic $Ca^{2+}$ activity following optogenetic stimulation of $PV_{IN}$ or $VGAT_{IN}$ Chr2 mice was impeded by miscellaneous activity but increased after systemic zolpidem administration.**

A) Number of active domains and area of active astrocytic $Ca^{2+}$ domains found before (control) and after 2-second optogenetic stimulation of anaesthetized $PV_{IN}$ or $VGAT_{IN}$ mice. No increase was found.

**B)** Astrocytic Ca2+ activity following optogenetic stimulation (both 2- and 15-second) measured as the area and number of active domains for $PV_{IN}$ stimulation in awake mice.

**C)** Same as (**B**) but for $VGAT_{IN}$.

**D)** Average number of active domains and area of active domains for both 2- and 15-second $PV_{IN}$ stimulation, as well as control recordings with no optogenetic stimulation. The top row shows no significant response under control conditions, whereas the bottom row shows the results after administration of the $GABA_A$ receptor allosteric modulator zolpidem. $^*p<0.05$.

**E)** Same as (**D**) but for $VGAT_{IN}$ stimulation.

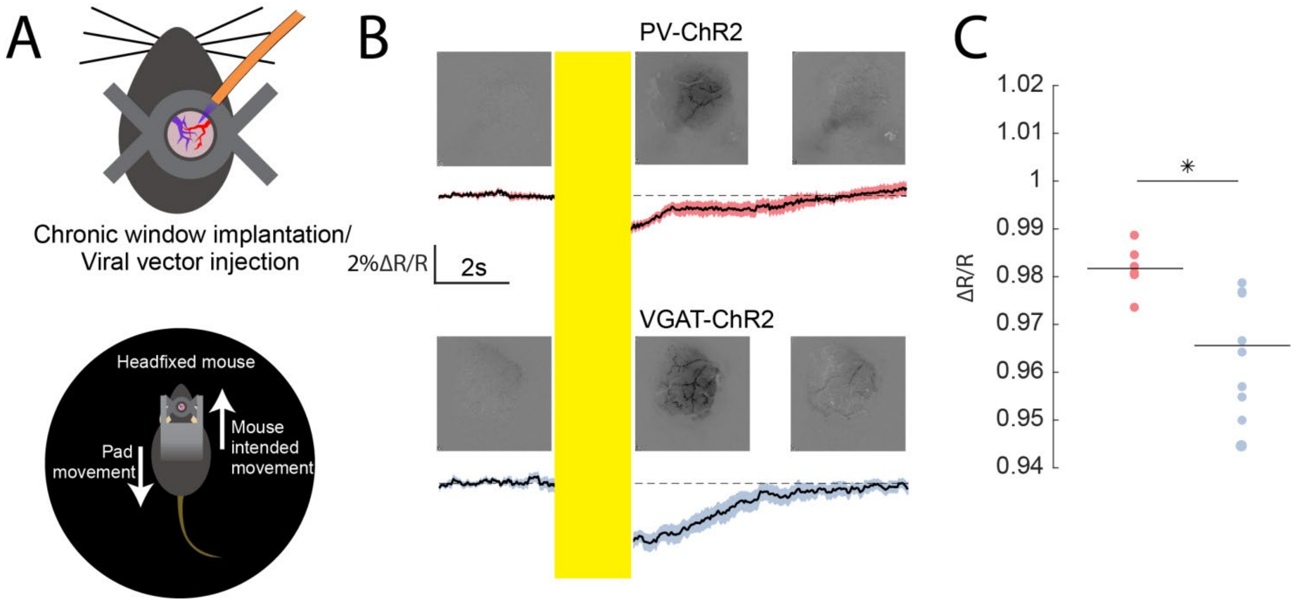


**Supplementary Figure S3:**

**$VGAT_{IN}$-based optogenetic stimulation induced larger cerebral blood flow (CBF) responses in awake mice than the $PV_{IN}$-based stimulation.**

A) Illustration of awake imaging and optogenetic stimulation setup.

**B)** IOS images from a single mouse and average IOS intensity traces for $PV_{IN}$ (red) and $VGAT_{IN}$ stimulation (blue) in response to 2-second optogenetic stimulation. Traces show normalized reflectance of the green light used for illumination. A decrease in reflectance indicates greater absorption of green light by haemoglobin in the blood, whereas a larger decrease in signal means a greater CBF response. Black lines show average traces and shading indicates the SEM.

**C)** Clear responses were observed in both strains, but the maximal negative amplitude of the reflectance and, thus, the amplitude of the CBF increase in $VGAT_{IN}$ stimulation was larger than the amplitude in $PV_{IN}$. *$p<0.05$.

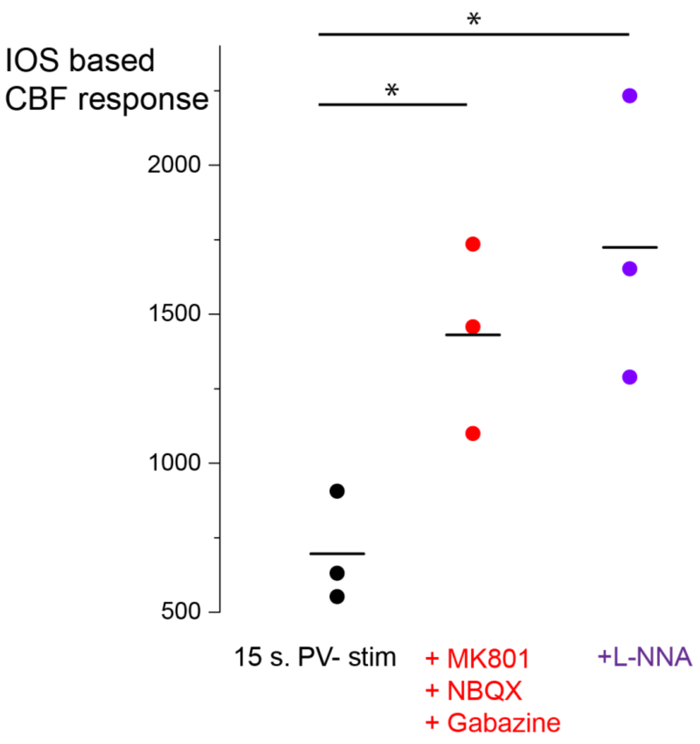


**Supplementary Figure S4:**

**Contribution of synaptic transmission, GABAA receptor, and NO to $PV_{IN}$-mediated cerebral blood flow (CBF) responses.**

Intense (15-s) optogenetic stimulation of $PV_{IN}$ mice induced a small CBF response in mice anaesthetized with α-chloralose. This response was increased after application of inhibitors of synaptic transmission (MK801 and NBQX) and GABAA receptor antagonist gabazine (red). The $PV_{IN}$-mediated CBF was not glutamate-dependent and gabazine counteracted the local inhibitory effect of the anaesthetics. Adding NOS inhibitor L-NNA did not significantly reduce the CBF response to PV interneuron activation (purple).
